# Physicochemical and Pharmacokinetic Properties Determining Drug Detection in Skin

**DOI:** 10.1101/2021.08.17.456720

**Authors:** Wout Bittremieux, Rohit S. Advani, Alan K. Jarmusch, Shaden Aguirre, Aileen Lu, Pieter C. Dorrestein, Shirley M. Tsunoda

## Abstract

Chemicals, including some systemically administered xenobiotics and their biotransformations, can be detected noninvasively using skin swabs and untargeted metabolomics analysis. We sought to understand the principal drivers that determine whether a drug taken orally or systemically is likely to be observed on the epidermis by using a random forest classifier to predict which drugs would be detected on the skin. A variety of molecular descriptors describing calculated properties of drugs, such as measures of volume, electronegativity, bond energy, and electrotopology, were used to train the classifier. The mean area under the ROC curve was 0.71 for predicting drug detection on the epidermis, and the SHapley Additive exPlanations model interpretation technique was used to determine the most relevant molecular descriptors. Based on the analysis of 2,561 FDA approved drugs, we predict that therapeutic drug classes such as nervous system drugs are more likely to be detected on the skin. Detecting drugs and other chemicals noninvasively on the skin using untargeted metabolomics could be a useful clinical advancement in therapeutic drug monitoring, adherence, and health status.

## Introduction

The skin provides a physical and chemical barrier to environmental insults and supports immunological function and thermoregulation. Additionally, the bacteria, viruses, and fungi that comprise the skin microbiome provide an essential function in protection against microbial pathogens and educating the immune system^1^. Traditionally, topical formulations of drugs are desired in certain medical conditions to either deliver drugs from the skin to the systemic circulation (e.g. transdermal scopolamine) or to deliver drugs locally to the skin and minimize the systemic toxicity of these drugs (e.g. topical corticosteroids). Interestingly, a recent study demonstrated systemic concentrations above the Food and Drug Administration (FDA) safety threshold of the sunscreen compounds avobenzone, oxybenzone, and octocrylene up to 21 days post administration, despite the widespread assumption that these commonly used topical products are considered “safe” ^2^.

An individual’s chemistry of their skin, including topically applied chemicals, such as avobenzone, octocrylene, as well as others found in soap, lotions, cosmetics, and anti-mosquito sprays and lotions can be detected in non-invasively obtained skin swab samples ^3^. We have recently demonstrated that systemically administered drugs, such as citalopram, diphenhydramine, and the N-acetyl metabolite of sulfamethoxazole, can be detected in skin swab samples of the hands, forearm, forehead, and axilla ^4^. Utilizing public untargeted human mass spectrometry metabolomics data and re-analysis of this data using the Global Natural Products Social Molecular Networking (GNPS)^5^ infrastructure, we achieved the detection of these compounds on the epidermis of patients that were prescribed these drugs, thus concluding that systemically administered drugs can be detected on the skin surface. Additionally, our recent study in healthy humans demonstrated a delayed time course between plasma and skin concentrations of diphenhydramine and its metabolites ranging from 1.5 to 10 hours (Panitchpakdi et al., in preparation).

The mechanism and pathways of chemicals and drugs moving from the systemic circulation to the epidermis are unknown. Additionally, not all xenobiotics can be detected on the skin. A notable example is the immunosuppressive drug tacrolimus, which was not detected in our skin swab samples ^4^. We sought to understand the physicochemical and pharmacokinetic properties that allow some systemically administered drugs to be detected on the epidermis and not others. Using existing skin swab data, we trained a random forest classifier to predict whether a compound will be observed on the epidermis.

## Methods

### Data Origin

No human subjects were recruited for this study and all data were assessed retrospectively (all data were anonymous and obtained from public metabolomics repositories). Publicly available tandem mass spectrometry (MS/MS) data on GNPS/MassIVE were accessed via ReDU (redu.ucsd.edu; ^6^ on March 24, 2019, as well as sample information (i.e. metadata) for the 30,626 files available at the time. Skin files were selected in the ReDU File Selector using the Uberon anatomy ontology ^7^ (Supplementary Table 1). This resulted in a list of 5,629 files, which were analyzed using MS/MS library searching (version 2.0; GNPS task ID: https://gnps.ucsd.edu/ProteoSAFe/status.jsp?task=53e265f8f6994f0196bf9bccd8d1b513). MS/MS library searching resulted in 175 drugs that were identified in the human skin files (level 2 annotation according to the Metabolomics Standards Initiative ^8^), filtered using a list of curated drugs and drug metabolites as they are recorded in the GNPS MS/MS reference libraries.

Duplicate annotations were removed and drugs available in topical formulations were excluded, resulting in a final list of 95 compounds. Based on the empirical measurement of these drugs or drug metabolites in publicly available MS data we presume that these 95 compounds are “positive” examples of drugs that appear on the epidermis. Additionally, we utilized the prescription records available in conjunction with data from a previous kidney transplant study ^4^ to define “negative” examples of drugs, i.e. drugs which were prescribed to individuals but were not observed in skin samples in that study. Skin samples were obtained from these individuals at two different visits for clinical evaluation. The subjects of that study were prescribed many (>5) medications simultaneously. Out of the 58 different medications in that study, 50 drugs were not detected in skin samples and offer “negative” examples for which we have experimental data. Negative examples will include both the lack of transport to the epidermis, but also the lack of detection due to sample preparation (e.g. some drugs might not be detected due to the chosen extraction conditions). The eight drugs or drug metabolites that were detected in skin swabs in that study are part of the presumed “positive” compounds. Further, these particular examples are supported with experimental data and matching prescription records (i.e. the drugs were detected in the subjects to whom they were prescribed).

Hence, 145 unique compounds were retained for the machine learning, of which 95 positive examples and 50 negative examples. The full list of compounds and information on whether they were observed on the epidermis or not is available in Supplementary Table 2.

### Machine Learning Epidermis Prediction

A random forest classifier was used to predict whether drugs are expected to be observed on the epidermis. First, Mordred ^9^, a cheminformatics software tool to efficiently compute a large variety of molecular descriptors, was used to generate molecular descriptors for all 145 compounds, such as calculated measures of volume, electronegativity, bond energy, and electrotopology (see Supplementary Table 3 for relevant descriptor examples). Molecular descriptors that were missing for one or more drugs were omitted, resulting in a feature table consisting of 929 unique descriptors per compound. Next, a classification pipeline was built to predict the probability of observing a drug on the epidermis. The classification pipeline consisted of preprocessing steps to remove irrelevant features and a random forest classifier. Preprocessing steps included removing all features whose variance was below 0.05 and removing one of the features for which their pairwise Pearson correlation exceeded 0.95. Next, a random forest classifier ^10^ using 1000 trees was trained to predict the epidermis probability.

Evaluation and hyperparameter tuning of the random forest was done using nested cross-validation. Two levels of stratified shuffle splitting consisting of 100 iterations of random splitting in 80% training data and 20% test data were performed. In the inner cross-validation loop, hyperparameter optimization was performed to determine the optimal depth of the trees in the random forest. Using grid search, tree depths between 5 and 9 (inclusive) were evaluated. The random forest classifier with the optimal tree depth was subsequently evaluated in the outer cross-validation loop. Trees with depth 8 were most frequently found to be optimal. For each split, the balanced accuracy, true positive rate, false positive rate, and precision were computed for both the training data and test data. Model performance was assessed based on the receiver operating characteristic (ROC) curve and precision–recall curve.

Important features for epidermis prediction were determined using SHAP (SHapley Additive exPlanations) ^11^, a model interpretability method founded in game theory. Briefly, SHAP explains machine learning predictions by using interpretable local models to approximate a complex black box model. Kernel SHAP was used to explore the trained classification pipeline. To determine the important features, 50 training samples determined by K-means clustering, with the cluster centroids weighted by the number of samples assigned to them, were used as the background dataset. To investigate the feature importances of individual compounds, if they were part of the training dataset, the random forest classifier with optimal hyperparameters was retrained using a leave-one-out strategy prior to SHAP analysis.

### FDA-Approved Drugs and Biotransformations

Drug names, SMILES representations, and ATC codes for 2,561 FDA approved drugs were retrieved from DrugBank (version 5.1.7) ^12^ on December 23rd, 2020. Mordred was used to generate the same features for these drugs as used during model training, and the probability of observing these drugs on the epidermis was determined using the trained classification pipeline. Additionally, potential biotransformation products of the drugs were generated using the BioTransformer tool ^13^. The human super transformer mode; which combines an Enzyme Commission-based transformer, a CYP450 (phase I) transformer, a phase II transformer, and a human gut microbial transformer; was used to predict potential biotransformation products after a single transformation step. This resulted in 23,693 putative biotransformation metabolites derived from the FDA approved drugs, for which similarly the probability of observing them on the epidermis was predicted using the trained classification pipeline.

### Code Availability

All analyses were performed in Python 3.8. RDKit (version 2020.09.3) ^14^ and Mordred (version 1.2.0) ^9^ were used to generate molecular descriptors. A GPU-accelerated version of the random forest algorithm, available as part of the cuML library (version 0.18.0) ^15^ was used in combination with Scikit-Learn (version 0.24.1) ^16^ for data preprocessing and model evaluation. SHAP (version 0.39.0) ^11^ was used to compute feature importances. BioTransformer (version 2.0.1) ^13^ was used to generate biotransformation products. Additionally, NumPy (version 1.20.1) ^17^, SciPy (version 1.6.0) ^18^, and Pandas (version 1.1.5) ^19^ were used for scientific computing, and matplotlib (version 3.3.4) ^20^ and Seaborn (version 0.11.1) ^21^ were used for visualization purposes.

All code is available at https://github.com/bittremieux/drugs_epidermis as open source under the permissive BSD license.

## Results

### Occurrence of Drugs on the Epidermis

Based on the rich metadata associated with the MS/MS data, we were able to select 5,629 publicly available MS/MS peak files that contain samples collected from human body sites. For our secondary analysis, we extracted 145 curated drugs to build a machine learning model to predict whether drugs occur on the epidermis. Additionally, based on the Uberon anatomy ontology ^7^, these drugs were mapped to the body site on which they were detected (Figure 1). The different rates of drug occurrence throughout the body suggest that there will be distinct detection of chemicals and xenobiotics in skin. As an example, our previous study showed that the N-acetyl metabolite of sulfamethoxazole was detected in armpit skin samples but not in other skin sites sampled such as forehead, palms, and forearm ^4^. More polar compounds may be more likely detected in more aqueous areas of the skin where sweat is more concentrated, such as the armpit.

**Figure 1:**
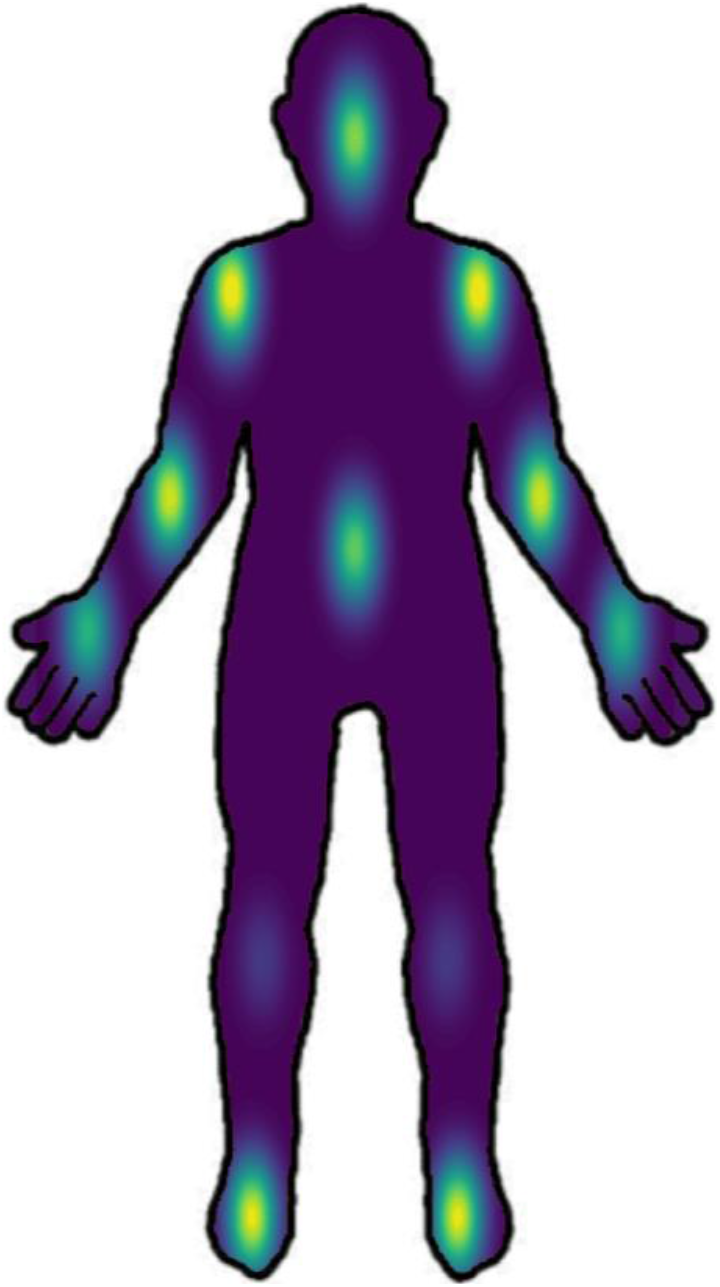
Body sites of the drugs found through spectral library searching. Body sites for the identified drugs were retrieved from the Uberon annotations specified in ReDU, and drug counts per body site were normalized by the total number of ReDU entries for each body site.

### Machine Learning to Predict Whether Drugs Occur on the Epidermis

Using a random forest classifier we were able to predict whether drugs will be observed on the epidermis with an area under the ROC curve (AUC) obtained during cross-validation of 0.71 ± 0.10 (Figure 2) and an area under the precision–recall curve of 0.82 ± 0.07 (Supplementary Figure 1). This performance indicates that machine learning can be used to successfully approximate the complex underlying biochemical processes leading to drugs being observed on the epidermis. As we were constrained by the limited availability of ground truth data in this study, we hypothesize that as more training data becomes available it will be possible to produce even more accurate machine learning models (Supplementary Figure 2).

**Figure 2.**
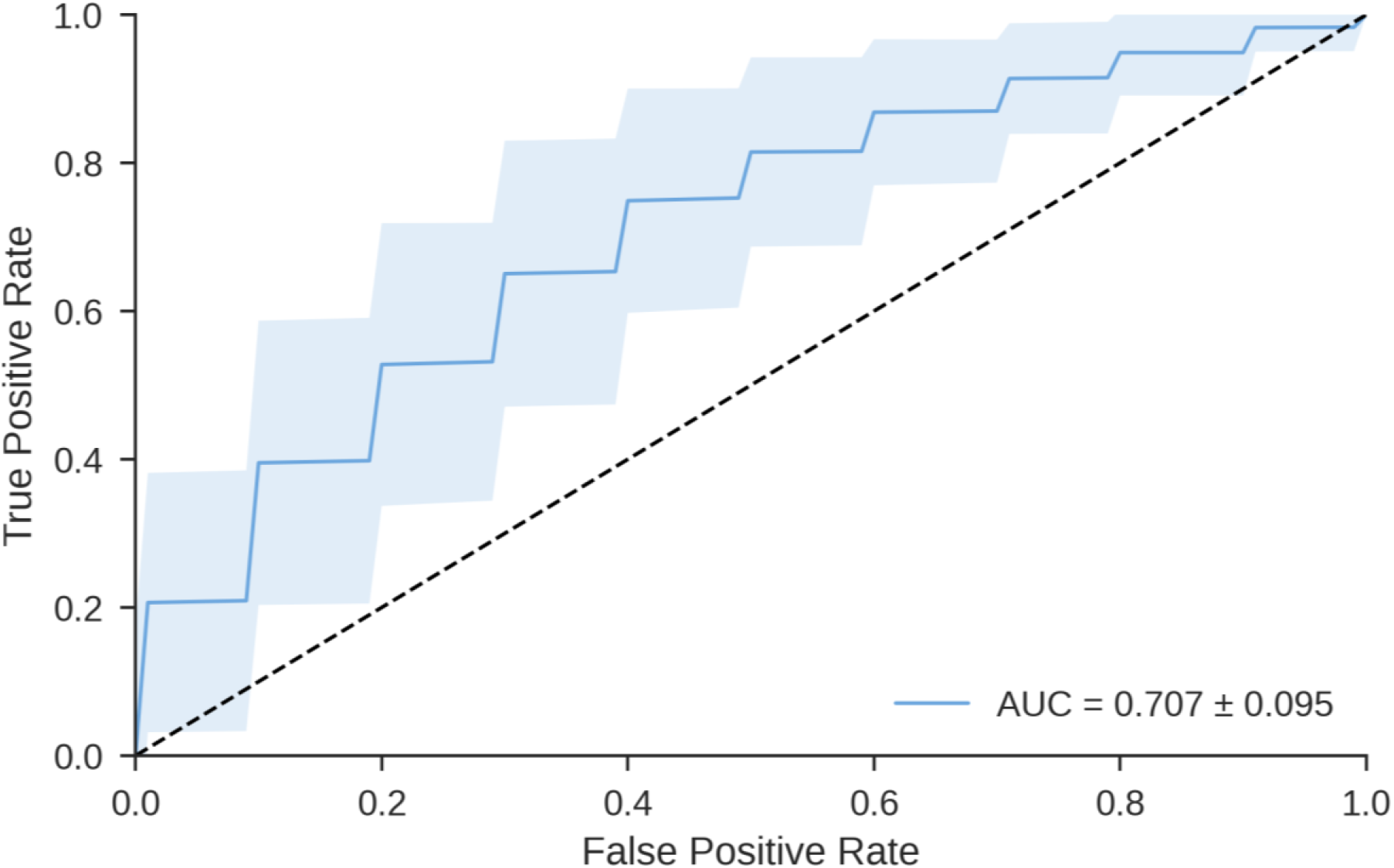
ROC curve indicating the performance of the random forest classifier to predict whether drugs can be observed on the epidermis. The curve is the mean ROC curve over 100 random stratified training (80% of the data) and test (20% of the data) splits. The standard deviation over the splits is indicated by the shaded area. The mean AUC is 0.707, with a standard deviation of 0.095.

We tried to gain insight into the molecules’ physical properties that results in drugs being present on the epidermis. The SHAP model interpretation technique was used to determine the most relevant features, consisting of molecular descriptors generated by Mordred, for the classifier performance (Figure 3). The top-ranked features are computed measures of volume (ATSC7v), electronegativity (PEOE_VSA1, PEOE_VSA9), bond energy (ATSC6d), and electrotopology (EState_VSA1). By investigating the SHAP values for individual features, we can derive that in general smaller compounds (Van der Waals volume) with a smaller bonding potential (electronegativity) are more likely to be observed on the epidermis. We can hypothesize that through heterogeneous biochemical processes such molecules diffuse faster and thus will be secreted to the epidermis.

**Figure 3.**
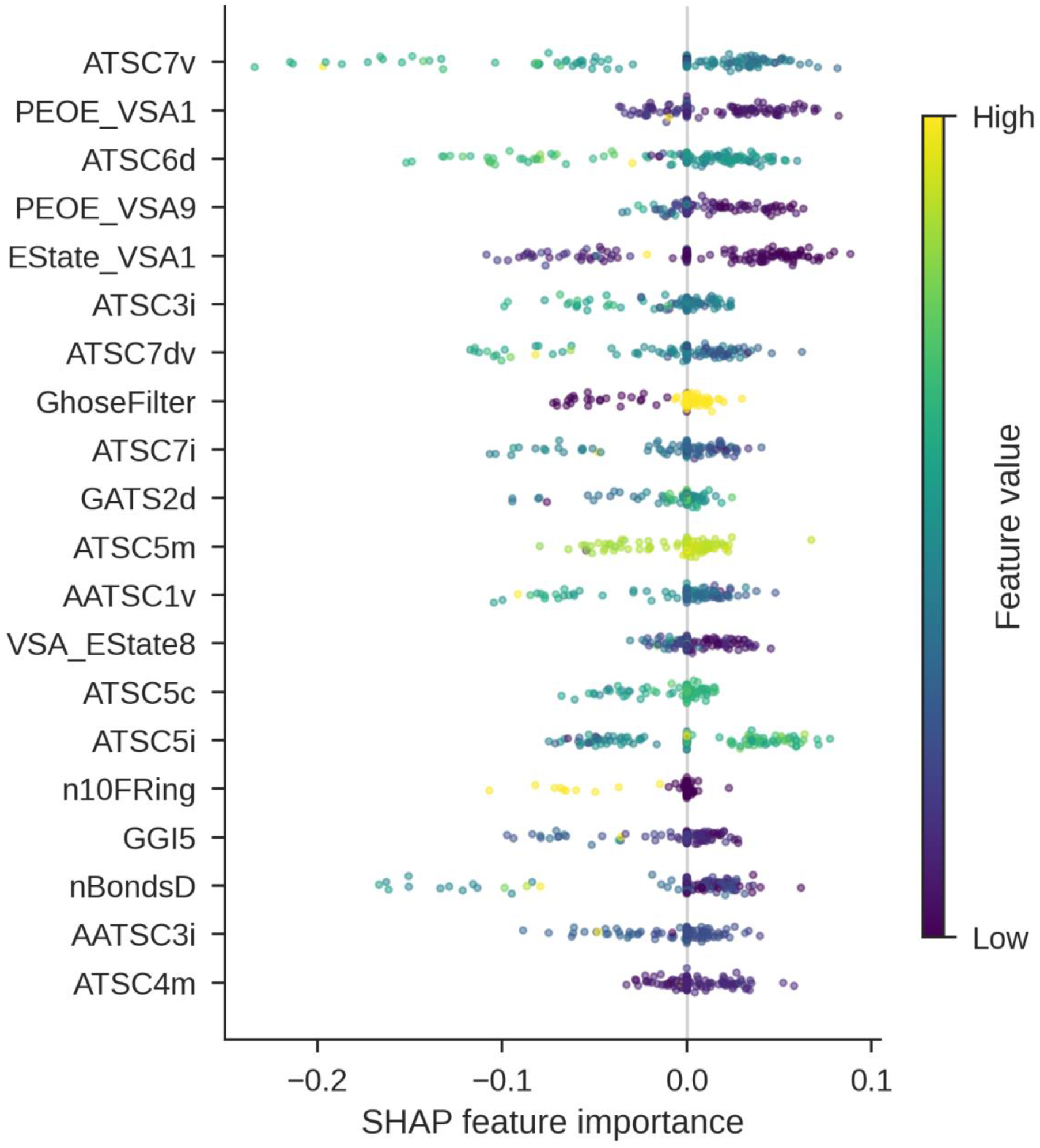
SHAP feature importances for the top 20 most important Mordred features from the random forest classifier for the 145 training compounds. A positive SHAP feature importance contributes to drugs predicted to appear on the epidermis, whereas a negative SHAP feature importance contributes to drugs predicted to not appear on the epidermis. The top-ranked features capture information about the volume, electronegativity, bond energy, and electrotopology of the molecules.

Additionally, SHAP can be used to interpret predictions for individual drugs. The antihistamine drug diphenhydramine was experimentally observed on the epidermis in a previous healthy human clinical study (Panitchpakdi et al. in preparation). Using a leave-one-out training strategy to not bias the classifier, it was also strongly predicted to be present on the epidermis (Figure 4A). The most relevant features contributing to this prediction are its low electrotopological state (EState_VSA1, VSA_EState8) and its low molecular connectivity and valence (Xch-7dv). In contrast, although the related compound diphenhydramine N-hexose is structurally similar, it is predicted to not appear on the epidermis (Figure 4B), in part because of its higher volume (ATSC7v), electrotopological state (EState_VSA1), polarizability (ATSC7p), and charge (GGI10). This is consistent with our experimental results (Panitchpakdi et al. in preparation). In a previous study ^4^, citalopram was detected in the skin samples of the only subject to which it was prescribed. This empirical observation is confirmed by the machine learning model (Figure 4C), as citalopram is very strongly predicted to be observed on the epidermis due to its low electrotopological state (EState_VSA1) and valence (ATSC7dv). Conversely, tacrolimus is very strongly predicted to not appear on the epidermis (Figure 4D), primarily due to its low ionization potential (ATSC8i), high number of double bonds (nBondsD), and its high number of Kier–Hall dssC atom types (motif “C(=[*])([*])[*]”) ^22^. This prediction matches its absence in the skin samples of 14 subjects who were prescribed tacrolimus ^4^. This analysis demonstrates how machine learning techniques can be used to obtain insights into the complex internal biochemical mechanisms that lead systemically administered drugs to be observed on the epidermis.

**Figure 4.**
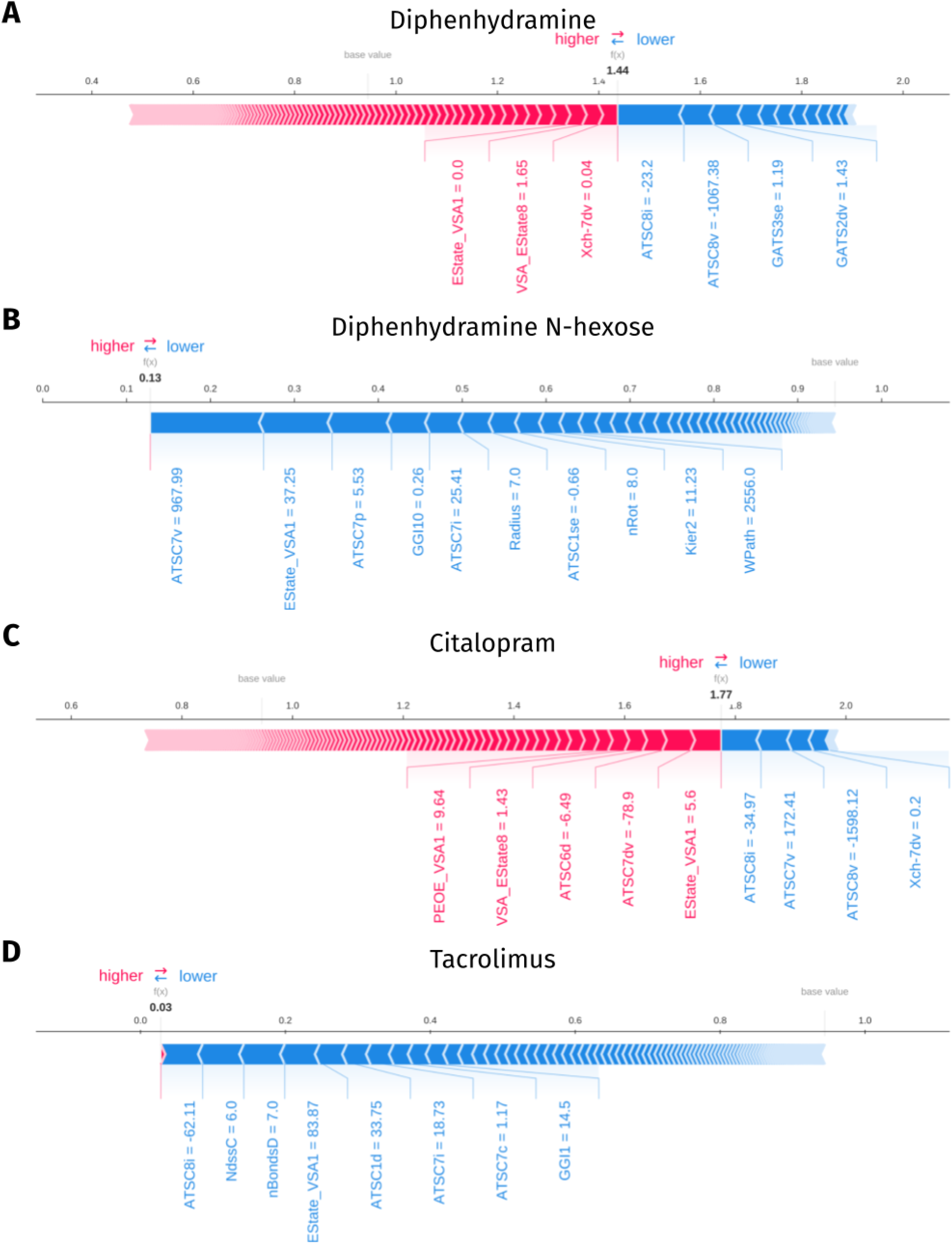
Force plots ^23^ of the SHAP values to interpret predictions of individual drugs. The most important features, their values, and the direction in which they contribute to the predictions (higher/red: observed, lower/blue: not observed) are displayed. Prediction values above the base value (indicated in grey) constitute positive predictions, values below the base value constitute negative predictions. **(A)** Diphenhydramine is predicted to be observed on the epidermis. **(B)** Diphenhydramine N-hexose is predicted to not be observed on the epidermis. **(C)** Citalopram is predicted to be observed on the epidermis. **(D)** Tacrolimus is predicted to not be observed on the epidermis.

To expand our knowledge of the variety of drugs that are likely to be observed on the epidermis beyond the training data consisting of 145 drugs, we retrieved 2,561 FDA approved drugs from DrugBank ^12^. Furthermore, we utilized BioTransformer ^13^ to predict potential biotransformation products of the FDA approved drugs, resulting in 23,693 putative biotransformation metabolites. These biotransformations include phase I metabolism products (e.g. Cytochrome P450), enzyme commission-based metabolism products, phase II metabolism products (e.g. Uridine 5’-diphospho-glucuronosyltransferase), and gut microbial transformation products, and they cover a number of different reaction types, including hydrolysis, oxidation and reduction, and conjugation.

The probability of observing both the FDA approved drugs and their potential biotransformation products was predicted using the trained random forest model. To investigate whether specific types of drugs were more likely to occur on the epidermis, we grouped the drugs and the corresponding biotransformation products using the Anatomical Therapeutic Chemical (ATC) Classification System (Figure 5). This indicates, for example, that hormonal preparations such as corticosteroids are least likely to be observed on the epidermis, while nervous system drugs such as analgesics, antiepileptics, antidepressants, and antipsychotics are more likely to be detected on skin.

**Figure 5.**
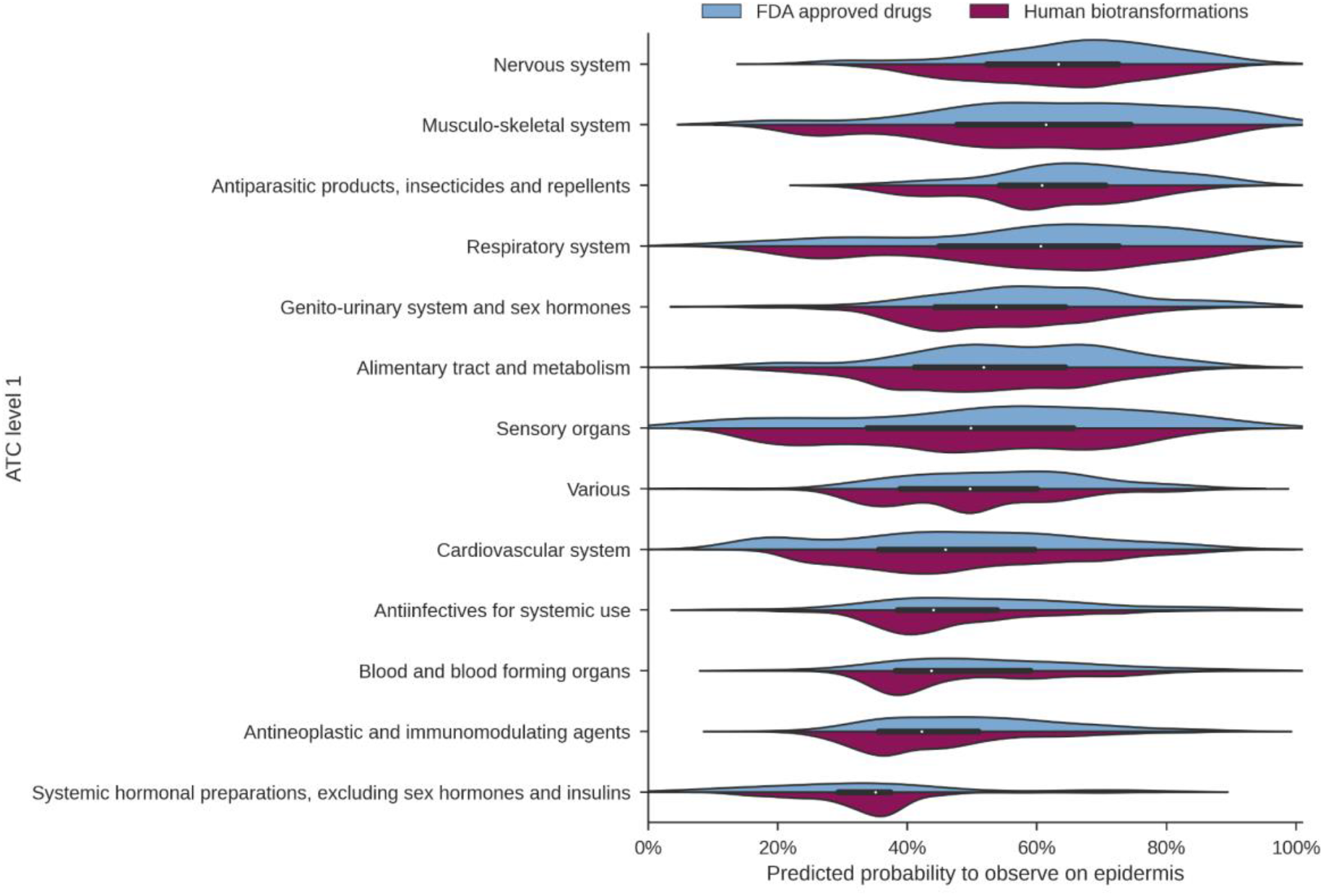
Prediction scores for 2,561 FDA approved drugs and their 23,693 biotransformations, subdivided by their drug class in the ATC classification system.

## Discussion

So far, little is known about which chemicals and drugs move from the systemic circulation to the epidermis. Here, we have demonstrated that machine learning can be used to gain insights into these complex processes for the first time. Using publicly available mass spectrometry data, we have trained a random forest model to predict whether drugs will occur on the epidermis. In addition to these secondary analysis results, the machine learning performance can be further improved upon by obtaining and incorporating more relevant experimental data, including both positive and negative examples of drugs and other xenobiotics that are commonly consumed and their status of being observed on the epidermis.

To obtain insights into the complex processes that underlie secretion of drugs to the epidermis, the SHAP model interpretability method was used to investigate which molecular descriptors are most relevant for prediction using the random forest. In general, we observe that smaller compounds with a smaller bonding potential are more likely to be observed on the epidermis.

Although further studies are needed to fully understand the underlying biochemical processes, we hypothesize that through heterogeneous mechanisms such molecules diffuse faster and thus will be secreted to the epidermis. Additionally, we used SHAP to investigate predictions for drugs with a known experimentally derived ground truth. This demonstrates how detailed and individualized insights for specific drugs can be obtained to explore whether they will appear on the epidermis or not.

Applying our random forest model to over 2,500 FDA approved drugs and their biotransformations gives insight into additional drugs and their metabolites that may be detected on the skin surface. For those drugs with low probability of skin detection, we hypothesize that either these drugs are fully processed within the body rather than secreted to the epidermis, or their physicochemical properties (e.g. high degree of lipophilicity) prevent access to the skin surface. Notably, median epidermis prediction values for the different ATC drug classes range from ∼35% to ∼60% of drugs in each category. There is a substantial variation in predicted probability; we speculate that this observation reflects that specific physicochemical properties of the drugs are the driver of this phenomena rather than the ATC class. Nevertheless, broad generalizations can be made; for example, steroid hormones were predicted to not be detected on the epidermis, which is consistent with our experimental data for budesonide, fludrocortisone, prednisone, and prednisolone; while amitriptyline, citalopram, cyclobenzaprine, escitalopram, gabapentin, ketamine, nortriptyline, and venlafaxine were detected in our data, consistent with our model prediction for nervous system drugs (Supplementary Table 2).

Our machine learning model is the first attempt to predict xenobiotic skin detection using physicochemical properties. There will likely be future iterations of this model as we advance our understanding of the complex processes governing molecular transport from the systemic circulation to the surface of the skin. The use of noninvasive skin swabs in clinical medicine could be a paradigm shift in how health and disease are monitored. Contemporary methods of blood draws and tissue biopsies are invasive and inconvenient for patients. In the future, we envision the use of noninvasive skin sampling to determine adherence of drugs, for therapeutic drug monitoring, extent of metabolism, and to assess organ and health status.

## Author Contributions

RSA, WB, AKJ, PCD, and SMT wrote the manuscript.

RSA, AKJ, and SMT designed the research.

RSA, WB, SA, AL, AKJ, and SMT performed the research.

RSA, WB, AKJ, and SMT analyzed the data.

RSA and WB contributed new reagents/analytical tools.

