## Supplemental Information for "Physicochemical and Pharmacokinetic Properties Determining Drug Detection in Skin"

### Supplementary Information

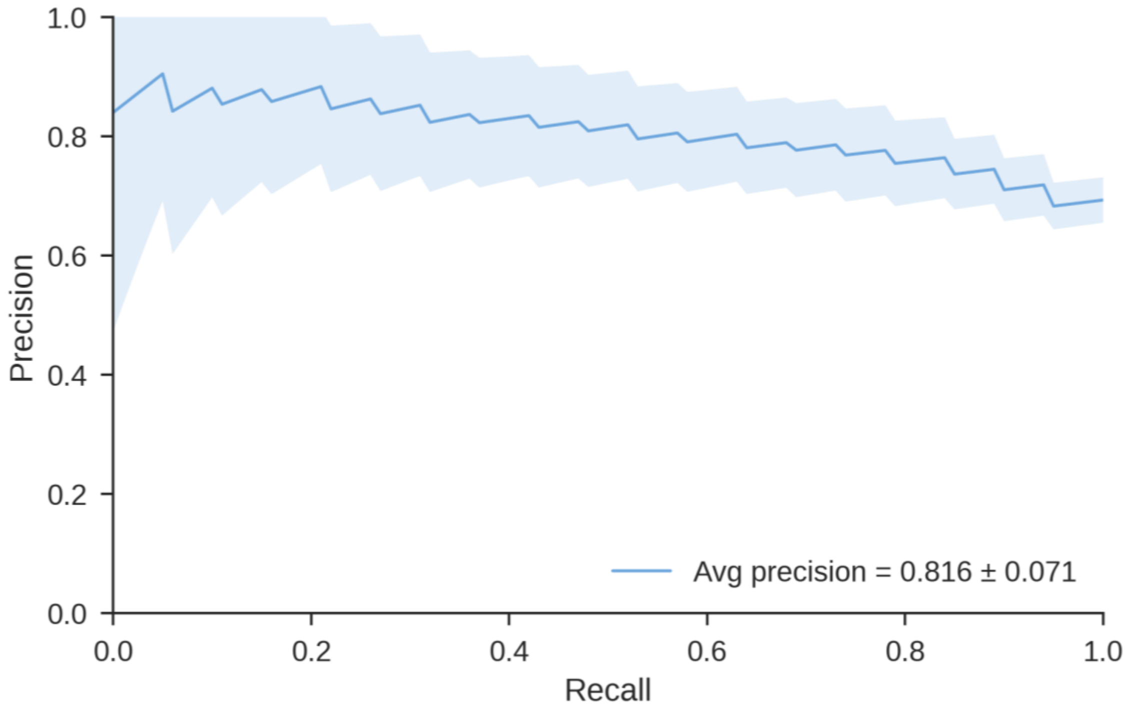


**Figure S1.** Precision–recall curve indicating the performance of the random forest classifier to predict whether drugs can be observed on the epidermis. The curve is the mean precision–recall curve over 100 random stratified training (80% of the data) and test (20% of the data) splits. The standard deviation over the splits is indicated by the shaded area. The mean area under the precision–recall curve is 0.816, with a standard deviation of 0.071.

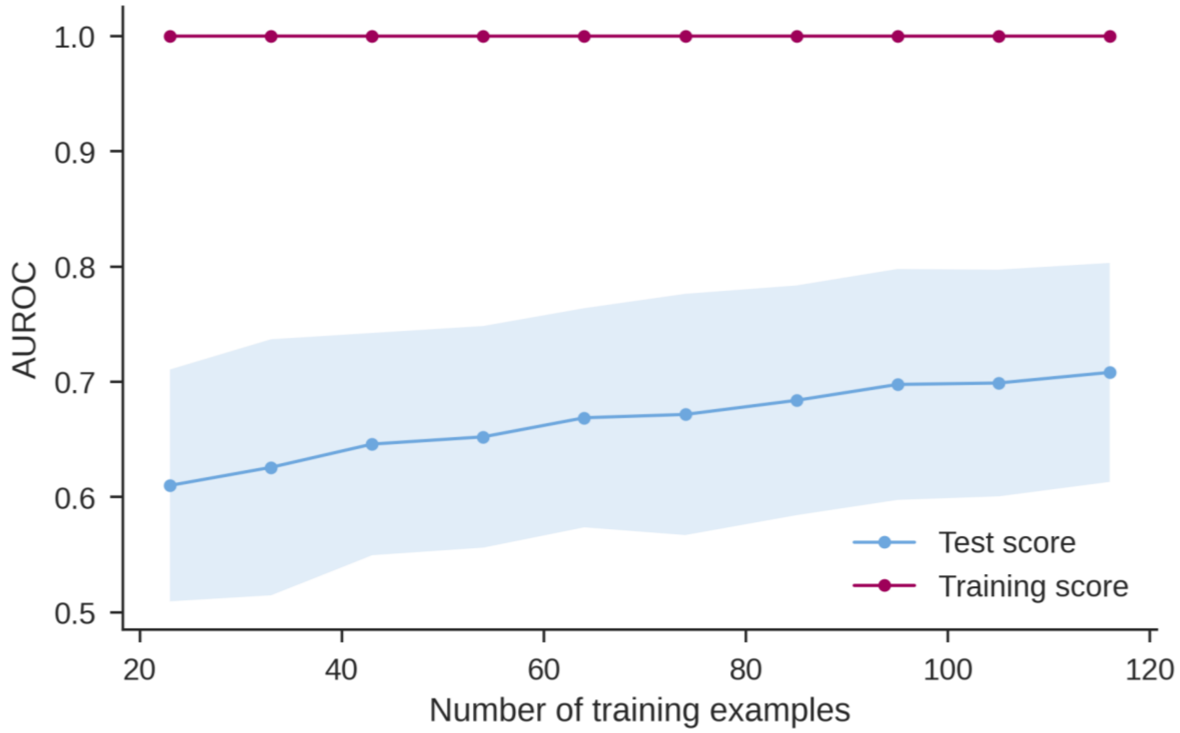


**Figure S2.** Learning curve to evaluate classification performance for an increasing number of training examples. Training and test performance was measured by the area under the ROC curve for 100 random stratified splits in 80% training and 20% test data, after which the number of training examples was subsampled to be between 20% and 100% of the available training data. The curves show the mean training and test scores, and the shaded areas show the standard deviation. The learning curve indicates that the performance increases as more training examples are available.

**Table S1:** Uberon ontology name and identifier used to select skin files available in ReDU.

| Name | Uberon identifier |
|---|---|
| arm skin | UBERON:0002427 |
| axilla skin | UBERON:0015474 |
| head or neck skin | UBERON:0012180 |
| skin of leg | UBERON:0001511 |
| skin of manus | UBERON:0001519 |
| skin of pes | UBERON:0001513 |
| skin of trunk | UBERON:0001085 |

**Table S2.** Training dataset with positive training samples that are observed on the epidermis and negative training samples that are not observed on the epidermis.

| **Compounds detected on epidermis** | **Compounds not detected on epidermis** |
|---|---|
| 11-Nor-9-carboxy-delta-9-tetrahydrocannabinol | Allopurinol |
| 2-Fluoromethcathinone | Amantadine |
| 2-Hydroxyibuprofen | Amlodipine |
| 4-Bromo-2,5-dimethoxymethamphetamine | Aspirin |
| 4-Fluoroisocathinone | Atenolol |
| 4-Hydroxycyclofenil | Atorvastatin |
| 6-Acetylmorphine | Baclofen |
| 6-beta-testosterone enanthate | Bisacodyl |
| Acetaminophen | Budesonide |
| Acetyl-sulfamethoxazole | Calcitriol |
| Acipimox | Cephalexin |
| Albendazole | Cholecalciferol |
| Ambroxol | Cinacalcet |
| Aminocaproic acid | Dapsone |
| Aminorex | Dutasteride |
| Amitriptyline | Emtricitabine |
| Amoxicillin | Ethenzamide |
| Ampicillin | Famotidine |
| Artemisinin | Fenofibrate |
| Azithromycin | Fludrocortisone |
| Bamethan | Furosemide |
| Benfluorex | Hydrochlorothiazide |
| Bestatin | Levothyroxine |
| Blebbistatin | Loperamide |

Caffeine
Cannabidiolic acid
Cefazolin
Chlorpheniramine
Citalopram
Clindamycin
Clotrimazole
Cyclobenzaprine
Cyclobenzaprine N-Oxide
Dacarbazine
Dehydroxy-amphotericin B
Dexchlorpheniramine
Dextromethorphan
Diclofenac
Dioctyl sulfosuccinate
Diphenhydramine
Disulfiram
Doxylamine
Ebastine
Escitalopram
Fenbendazole
Finasteride
Fluconazole
Gabapentin
Gestiron
Guaifenesin
Ibudilast

Loratadine
Lorazepam
Methenamine
Metoprolol
Montelukast
Nifedipine
Omeprazole
Oxycodone
Pramipexole
Prednisolone
Prednisone
Pregnenedione
Prochlorperazine
Promethazine
Propranolol
Raltegravir
Rosuvastatin
Simvastatin
Sodium Bicarbonate
Tacrolimus
Tamsulosin
Tenofovir Alafenamide
Tramadol
Valganciclovir
Vancomycin
Zolpidem

Ibuprofen

Indomethacin

Isoconazole

Isoprenaline

Ketamine

Ketoconazole

Linezolid

Miconazole

Morphine

Mycophenolate mofetil

Mycophenolic acid

N-Acetylsulfamethoxazole

N-Desmethylcyclobenzaprine

N-Desmethyltapentadol

N-Methylephedrine

Nadifloxacin

Naproxen

Niclosamide

Nifenazone

Norketamine

Norphenylephrine

Nortriptyline

Nystatin

Oxandrolone

Prasterone

Procaine

Proguanil

| |
|---|
| Ranitidine |
| Rivanicline |
| Sulfacetamide |
| Sulfachloropyridazine |
| Sulfadiazine |
| Sulfadimethoxine |
| Sulfadoxine |
| Sulfafurazole |
| Sulfamethizole |
| Sulfamethoxazole |
| Thiabendazole |
| Tolnaftate |
| Trimethoprim |
| Valsartan |
| Venlafaxine |
| Venlafaxine N-Oxide |
| Warfarin |

**Table S3.** List of the top-ranked Mordred features determined by SHAP. The full list of descriptors computed by Mordred is available at https://mordred-descriptor.github.io/documentation/master/descriptors.html.

| **Mordred descriptor** | **Definition** |
|---|---|
| ATSC7v | centered Moreau-Broto autocorrelation of lag 7 weighted by Van der Waals volume |
| PEOE_VSA1 | MOE charge Van der Waals surface area descriptor 1 (-inf < x < -0.30) |
| ATSC6d | autocorrelation of lag 6 weighted by sigma electrons |
| PEOE_VSA9 | MOE charge Van der Waals surface area descriptor 9 (0.05 ≤ x < 0.10) |
| EState_VSA1 | Electrotopological state Van der Waals surface area descriptor 1 (-inf < x < -0.39) |
| ATSC3i | centered Moreau-Broto autocorrelation of lag 3 weighted by ionization potential |
| ATSC7dv | centered Moreau-Broto autocorrelation of lag 7 weighted by valence electrons |
| GhoseFilter | Ghose filter |
| ATSC7i | centered Moreau-Broto autocorrelation of lag 7 weighted by ionization potential |
| GATS2d | Geary’s coefficient of lag 2 weighted by sigma electrons |
| ATSC5m | centered Moreau-Broto autocorrelation of lag 5 weighted by mass |
| AATSC1v | averaged and centered Moreau-Broto autocorrelation of lag 1 weighted by Van der Waals volume |
| VSA_Estate8 | Van der Waals surface area electrotopological state descriptor 8 (6.45 ≤ x < 7.00) |
| ATSC5c | centered Moreau-Broto autocorrelation of lag 5 weighted by Gasteiger charge |
| ATSC5i | centered Moreau-Broto autocorrelation of lag 5 weighted by ionization potential |
| n10FRing | 10-membered fused ring count |
| GGI5 | 5-ordered raw topological charge |
| nBondsD | number of double bonds in non-kekulized structure |
| AATSC3i | averaged and centered Moreau-Broto autocorrelation of lag 3 weighted by ionization potential |
| ATSC4m | centered Moreau-Broto autocorrelation of lag 4 weighted by mass |
